# *Exaiptasia diaphana* from the Great Barrier Reef: a valuable resource for coral symbiosis research

**DOI:** 10.1101/775510

**Authors:** Ashley M. Dungan, Leon Hartman, Giada Tortorelli, Roy Belderok, Annika M. Lamb, Lynn Pisan, Geoffrey I. McFadden, Linda L. Blackall, Madeleine J. H. van Oppen

## Abstract

The sea anemone, *Exaiptasia diaphana*, commonly known as *Exaiptasia pallida* or *Aiptasia pallida*, has become increasingly popular as a model for cnidarian-microbiome symbiosis studies due to its relatively rapid growth, ability to reproduce sexually and asexually, and symbiosis with diverse prokaryotes and the same microalgal symbionts (family Symbiodiniaceae) as its coral relatives. Clonal *E. diaphana* strains from Hawaii, the Atlantic Ocean, and Red Sea are now established for use in research. Here, we introduce Great Barrier Reef (GBR)-sourced *E. diaphana* strains as additions to the model repertoire. Sequencing of the 18S rRNA gene confirmed the anemones to be *E. diaphana* while genome-wide single nucleotide polymorphism analysis revealed four distinct genotypes. Based on *Exaiptasia*-specific inter-simple sequence repeat (ISSR)-derived sequence characterized amplified region (SCAR) marker and gene loci data, these four *E. diaphana* genotypes are distributed across several divergent phylogenetic clades with no clear phylogeographical pattern. The GBR *E. diaphana* genotypes comprised three females and one male, which all host *Breviolum minutum* as their homologous Symbiodiniaceae endosymbiont. When acclimating to an increase in light levels from 12 to 28 μmol photons m^-2^ s^-1^, the genotypes exhibited significant variation in maximum quantum yield of Symbiodiniaceae photosystem II and Symbiodiniaceae cell density. The comparatively high levels of physiological and genetic variability among GBR anemone genotypes makes these animals representative of global *E. diaphana* diversity and thus excellent model organisms. The addition of these GBR strains to the worldwide *E. diaphana* collection will contribute to cnidarian symbiosis research, particularly in relation to the climate resilience of coral reefs.

## 1. Introduction

### 1.1 Coral Reefs

The Great Barrier Reef (GBR) contains abundant and diverse biota, including more than 300 species of stony corals (Fabricius et al. 2005), making it one of the most unique and complex ecosystems in the world. In addition to its tremendous environmental importance, the GBR’s social and economic value is estimated at $A56 billion, supporting 64,000 jobs and injecting $A6.4 billion into the Australian economy annually (O’Mahony et al. 2017).

Coral reef waters are typically oligotrophic, but stony corals thrive in this environment and secrete the calcium carbonate skeletons that create the physical structure of the reef. Corals achieve this through their symbiosis with single-celled algae of the family Symbiodiniaceae that reside within the animal cells and provide the host with most of its energy (Muscatine and Porter 1977). Additional support is provided by communities of prokaryotes, and possibly by other microbes such as viruses, fungi and endolithic algae living in close association with coral. This entity, comprising the host and its microbial partners, is termed the holobiont (Rohwer et al. 2002). During periods of extreme thermal stress, the coral-Symbiodiniaceae relationship breaks down. Stress-induced cellular damage creates a state of physiological dysfunction, which leads to separation of the partners and, potentially, death of the coral animal. This process, ‘coral bleaching’, is a substantial contributor to coral cover loss globally (Baird et al. 2009; De’ath et al. 2012; Eakin et al. 2016; Hughes et al. 2017; Hughes et al. 2018).

### 1.2 Exaiptasia diaphana

Model systems are widely used to explore research questions where experimentation on the system of interest has limited feasibility or ethical constraints. Established model systems, such as *Drosophila melanogaster* and *Caenorhabditis elegans*, have played crucial roles in the progress of understanding organismal function and evolution in the past 50 years (Davis 2004). The coral model *Exaiptasia diaphana* was formally proposed by Weis et al. (2008) as a useful system to study cnidarian endosymbiosis and has since achieved widespread and successful use.

*E. diaphana* is a small sea anemone (≤60 mm long) found globally within temperate and tropical marine shallow-water environments (Grajales and Rodriguez 2014). Originally positioned taxonomically within the genus *Aiptasia, E. diaphana* and twelve other *Aiptasia* species were combined into a new genus, *Exaiptasia* (Grajales and Rodriguez 2014). Although *Exaiptasia pallida* was proposed as the taxonomic name for the twelve synonymized species, the International Commission on Zoological Nomenclature (ICZN) ruled against this because the species epithet *diaphana* (Rapp 1829) predated *pallida* (Verrill 1864) and therefore had precedence according to the Principle of Priority (ICZN 2017).

*E. diaphana* was first used to study cellular regeneration (Blanquet and Lenhoff 1966), with its systematic use in the study of cnidarian-algal symbioses dating back to 1976 when the regulation of *in hospite* Symbiodiniaceae density was explored (Steele 1976). Since then, studies using *E. diaphana* have focused on the onset, maintenance, and disruption of symbiosis with Symbiodiniaceae (Belda-Baillie et al. 2002; Fransolet et al. 2014; Bucher et al. 2016; Hillyer et al. 2017; Cziesielski et al. 2018). *E. diaphana* has also been used in studies of toxicity (Duckworth et al. 2017; Howe et al. 2017), ocean acidification (Hoadley et al. 2015), disease and probiotics (Alagely et al. 2011), and cnidarian development (Chen et al. 2008; Grawunder et al. 2015; Carlisle et al. 2017). Key differences between corals and *E. diaphana* are the absence of a calcium carbonate skeleton, the constant production of asexual propagates, and the greater ability to survive bleaching events in the latter. These features allow researchers to use *E. diaphana* to investigate cellular processes that would otherwise be difficult with corals, such as those that require survival post-bleaching to track re-establishment of eukaryotic and prokaryotic symbionts. Further, adult anemones can be fully bleached without eliciting mortality, which is often difficult for corals, thus providing a system for algal reinfection studies independent of sexual reproduction and aposymbiotic larvae.

Three strains of *E. diaphana* currently dominate the research field as models for coral research. H2 (female) was originally collected from Coconut Island, Hawaii, USA (Xiang et al. 2013), CC7 (male) from the South Atlantic Ocean off North Carolina, USA (Sunagawa et al. 2009) and RS (female, pers.comm., but see (Schlesinger et al. 2010)) collected from the Red Sea at Al Lith, Saudia Arabia (Cziesielski et al. 2018). The majority of *E. diaphana* resources have been developed from the clonal line, CC7, including reproductive studies (Grawunder et al. 2015), transcriptomes (Sunagawa et al. 2009; Lehnert et al. 2012; Lehnert et al. 2014) and an anemone genome sequence (Baumgarten et al. 2015). Prior work evaluating multiple *E. diaphana* strains has shown variation in sexual reproduction (Grawunder et al. 2015), Symbiodiniaceae specificity (Thornhill et al. 2013; Grawunder et al. 2015), and resistance to thermal stress (Bellis and Denver 2017; Cziesielski et al. 2018). However, Australian contributions to these efforts have been hampered as researchers have not had access to the established strains as import permits for alien species are difficult to secure.

The aim of the present work is to add GBR representatives to the existing suite of *E. diaphana* model strains. We describe the establishment and maintenance of four genotypes, including their provenance, taxonomy, maintenance, histology, physiology, and Symbiodiniaceae symbiont type. The results highlight characteristics that make GBR-sourced *E. diaphana* a valuable extension to this coral model, such as the physiological and genetic variability between genotypes, which is more representative of a global population. This foundation information will be useful to international researchers interested in using the GBR-sourced animals in their research and will give Australian researchers access to this valuable model.

## 2. Methods

### 2.1 Anemone acquisition and husbandry

Several pieces of coral rubble bearing anemones were obtained from holding tanks in the National Sea Simulator (SeaSim) at the Australian Institute of Marine Science and sent to Swinburne University of Technology, Melbourne (SUT) in late 2014. Additional anemones from the SeaSim were sent to the Marine Microbial Symbiont Facility (MMSF) at the University of Melbourne (UoM) in early 2016. Most material in the SeaSim tanks originates from the central GBR; therefore, this is the likely origin of the anemones. The SUT and UoM populations were consolidated at MMSF in early 2017 where they were segregated according to single polynucleotide polymorphism (SNP) analysis groupings (see below). Anemones were grown in 4 L polycarbonate tanks in reverse osmosis (RO) water reconstituted Red Sea Salt(tm) (RSS) at ∼34 parts per thousand (ppt), and incubated without aeration or water flow at 26°C under lighting of 12-20 μmol photons m^−2^ s^−1^ (light emitting diode - LED white light array) on a 12h:12h light:dark cycle in a walk-in incubator. Anemones were fed *ad libitum* with freshly hatched *Artemia salina* (brine shrimp, Salt Creek, UT, USA) nauplii twice weekly. Tanks were cleaned each week after feeding by loosening algal debris with water pressure applied through disposable plastic pipettes, removing algal biomass, and complete water changes. When cleaning, ∼25% of the anemones were cut into 2-6 fragments to promote population expansion through regeneration of the tissue fragments into whole anemones. Every third week, all anemones were transferred to clean tanks.

### 2.2 Anemone identity and genotyping

Anemone identity was determined by Sanger sequencing of the 18S rRNA gene. Genomic DNA (gDNA) was extracted from six whole anemones following the protocol described by Wilson et al. (2002), modified with a 15 min incubation in 180 μM lysozyme, and 30 s bead beating at 30 Hz (Qiagen Tissue-Lyser II) with 100 mg of sterile glass beads (Sigma G8772). The 18S rRNA genes were PCR amplified from all anemone samples using external Actiniaria-specific 18S rRNA gene primers 18S_NA, 5’-TAAGCACTTGTCTGTGAAACTGCGA-3’ and 18S_NB, 5’-TAAGCACTTGT CTGTGAAACTGCGA-3’ (Grajales and Rodriguez 2016) with 0.5 U 2x Mango Mix (Bioline), 2 μL of DNA template, 0.2 μM of each primer, and nuclease-free water up to 25 μL. PCR conditions consisted of initial denaturation at 94°C for 5 min, 35 cycles of 94°C for 45 s, 55°C for 45 s, and 72°C for 45 s followed by a final extension at 72°C for 5 min. PCR products were purified with an ISOLATE II PCR and Gel Kit (Bioline, BIO-52059) according to the manufacturers guidelines, and supplied to the Australian Genome Research Facility (AGRF) for Sanger sequencing with the external primers and four internal primers: 18S_NL, 5’-AACAGCCCGGTCAGTAACACG-3’, 18S_NC 5’-AATAACAATACAGGGCTTTTCTAAGTC-3’, 18S_NY 5’-GCCTTCCTGACTTTGGTTGAA-3’, and 18S_NO 5’-AGTGTTATTGGATGACCTCTTTGGC-3’. The raw 18S reads were aligned in Geneious (v 10.0.4) (Kearse et al. 2012) to produce the near-complete 18S rRNA gene sequence, which was evaluated by BLASTn (Altschul et al. 1990) to identify the anemones.

For genotyping, DNA was extracted as described above from 23 whole anemones or their tentacles and sent to Diversity Arrays Technology Pty Ltd (Canberra, Australia) for DArT next-generation sequencing (DArTseq). DArTseq combines complexity reduction and next generation sequencing to generate genomic data with a balance of genome-wide representation and coverage (Cruz et al. 2013). Complexity reduction was achieved by using restriction endonucleases to target low-copy DNA regions. These regions were then sequenced using Illumina HiSeq2500 (Illumina, USA) with an average read depth exceeding 20x. The data were processed by DArT Pty Ltd to remove poor quality sequences and to ensure reliable assignment of sequences to samples. DArTsoft14 was then used to identify SNPs at each locus as homozygous reference, homozygous alternate or heterozygous for each individual (Melville et al. 2017). Monomorphic loci, loci with <100% reproducibility or missing values were removed in the R package, dartR, to improve the quality of and reduce linkage within the dataset; this reduced the dataset from 8288 loci to 1743 loci (R Core Team 2013; Gruber et al. 2017).

Euclidean distances between individual anemones, based on differences in the allele frequencies at each of the SNP loci, were calculated from the reduced SNP dataset in dartR, then viewed as a histogram and printed into a matrix in RStudio. Although the genetic distance between individuals of the same genotype should be zero, small differences may occur due to sequencing errors and somatic mutations. The genetic distance between individuals of different genotypes will be larger than that within individuals. Therefore, the genetic distances among individuals from several genotypes should form a bi- or multi-modal distribution; one peak with a relatively small mean represents genetic distances between pairs of individuals of the same genotype, another represents inter-genotypic distances with a larger mean. The inter-genotypic distribution can be multi-modal because different pairs of genotypes can differ by different amounts. Note that two samples from the same individual were genotyped to determine methodological error rates and verify the baseline for clonality. A principal coordinates analysis was performed and plotted in dartR to visualize the genotype assignments.

To compare the phylogenetic relationship of the GBR-sourced anemones with the previously described clonal lines, we used a set of four *Exaiptasia*-specific inter-simple sequence repeat (ISSR)-derived sequence characterized amplified region (SCAR) markers developed by Thornhill et al. (2013) and an additional six *Exaiptasia*-specific gene loci (Bellis and Denver 2017). gDNA was extracted from five animals of each genotype, as described above, and amplified with each of the four SCAR marker (3, 4, 5, and 7) and gene primer pairs (Thornhill et al. 2013; Bellis and Denver 2017). PCR solutions for each marker contained 0.5U MyTaq HS Mix polymerase (Bioline), 1 μL of DNA template, 0.4 μM of each primer, and nuclease-free water up to 25 μL. Thermocycling consisted of an initial denaturation at 94°C for 1.5 min, 35 cycles of 94°C for 1 min, 56°C for 1 min, and 72 °C for 1.5 min followed by a final extension at 72°C for 5 min. Amplified products were purified and sequenced in forward and reverse directions at AGRF. Sequences were aligned and edited in Geneious version 2019.2. Substantial non-specific binding of the SCAR marker 7 primers generated unusable sequence data. Further, the forward and reverse sequences from two anemone genotypes that were heterozygous for two or more indels at different points in the sequences of SCAR markers 3 (anemone AIMS1) and 4 (anemones AIMS2-4) could not be aligned and this prohibited the inclusion of those loci from analyses. Therefore, only sequence data from SCAR marker 5 was used for phylogenetic analyses. For the six *Exaiptasia*-specific gene loci, only AIPGENE19577, Atrophin-1-interacting protein 1 (AIP1) contained heterozygous indels or non-specific binding of the forward primer.

For each genotype, the SCAR marker 5 sequences and each of the six *Exaiptasia*-specific gene loci of the clonal replicates were aligned and a consensus sequence was generated. For the SCAR marker, the consensus sequences were aligned with twelve reference sequences from Thornhill et al. (2013) and three experimental anemone sequences (Grawunder et al. 2015), while the *Exaiptasia*-specific gene sequences were aligned with five reference sequences from Bellis and Denver (2017). Each alignment was used to create a phylogenetic tree using the Maximum Likelihood method and General Time Reversible model (Nei and Kumar 2000) in MEGA X (Kumar et al. 2018). Topology, branch length, and substitution rate were optimized, and branch support was estimated by bootstrap analysis of 1000 iterations.

### 2.3 GBR anemone gender determination

Over a period of two months, several anemone individuals from each genotype were reared to a pedal disk diameter of ∼7 mm. One animal per genotype was anesthetized in a 1:1 solution of 0.37 M magnesium chloride and filter-sterilized RSS (fRSS) for 30 min, dissected and viewed under a stereo microscope (Leica MZ8) to confirm the presence of gonads. When gonads were observed, one animal per genotype was anesthetized as described above, fixed in 4% paraformaldehyde in 1x PBS (pH 7.4) overnight, washed three times in 1x PBS and processed by the Biomedical Histology Facility of The University of Melbourne. Samples were embedded in paraffin and 7 μm transverse sections were cut (see supplemental information Online Resource 1 for histological processing). Slides were stained with hematoxylin and eosin (H&E) and viewed under a compound microscope (Leica DM6000B).

### 2.4 Symbiodiniaceae

The ITS2 region of Symbiodiniaceae gDNA was analyzed by metabarcoding to determine the Symbiodiniaceae species associated with the GBR-sourced anemones. gDNA extracted from three animals of each genotype, as described above, was amplified in triplicate with the ITS-Dino (forward; 5’-<u>TCGTCGGCAGCGTCAGATGTGTATAAGAGACAG</u>GTGAATTGCAGAACTCCGTG-3’) (Pochon et al. 2001) and its2rev2 (reverse; 5’-<u>GTCTCGTGGGCTCGGAGATGTGTATAAGAGACAG</u>CCTCCGCTTACTTATATGCTT-3’) (Stat et al. 2009) primers modified with <u>Illumina adapters</u>. PCRs included 0.5 U MyTaq HS Mix polymerase (Bioline), 1 μL of DNA template, 0.25 μM of each primer, and nuclease-free water up to 20 μL. Thermocycling consisted of an initial denaturation at 95.0 °C for 3 min, 35 cycles at 95.0 °C, 55.0 °C and 72.0 °C for 15 sec each, and 1 cycle at 72 °C for 3 min. Library preparation on pooled triplicates and Illumina MiSeq sequencing (2×250 bp) was performed by the Ramaciotti Centre for Genomics at the University of New South Wales, Sydney. Raw, demultiplexed MiSeq read-pairs were joined in QIIME2 v2018.4.0 (Bolyen et al. 2018). Denoising, chimera checking, and trimming was performed in DADA2 (Callahan et al. 2016). The remaining sequences were clustered into operational taxonomic units (OTUs) at 99% sequence similarity using closed-reference OTU picking in vsearch (Rognes et al. 2016). A taxonomic database adapted from Arif et al. (2014) was used to seed the OTU clusters and for taxonomic classification.

### 2.5 Anemone and Symbiodiniaceae physiological properties

Physiological assessments were performed on anemones maintained in a healthy, unbleached state for over two years. Three of the four anemone genotypes named AIMS2, AIMS3, and AIMS4, with oral disk diameters of 3-4 mm were transferred from the original holding tanks in the walk-in incubator and randomly distributed between three replicate (by genotype) 250 mL glass culture containers containing RSS. Anemones of this size were chosen as they were considered sexually immature (Muller-Parker et al. 1990; Grawunder et al. 2015) and therefore variability in sexual development or gonadal reserves were unlikely to influence the results. AIMS1 was excluded from long-term assessment as it had poor survival at densities required for the analyses.

Glass culture containers were transferred from the walk-in incubator to experimental growth chambers (Taiwan HiPoint Corporation model 740FHC) fitted with red, white, and infrared LED lights. Initial light levels were set at 12 μmol photons m^-2^ s^-1^ (HiPoint HR-350 LED meter) to correspond with the walk-in incubator conditions and ramped up to 28 μmol photons m^-2^ s^-1^ over a period of 72 h on a 12 h:12 h light:dark cycle to approximate experimental conditions reported in *E. diaphana* literature (Online Resource 2; Fransolet et al. 2014; Hoadley and Warner 2017). While the commonly used *E. diaphana* strains (CC7, H2, RS) are often maintained at >50 μmol photons m^−2^ s^−1^ (pers. comm.), the GBR strains appeared healthier, with extended bodies and open tentacles, at lower light intensities. The anemones were maintained in RSS, fed *A. salina* nauplii and cleaned as described above. The pH, salinity and temperature of the water and applied light levels were monitored thrice weekly and were stable over time: 8.14±0.02, 35.0±0.04 ppt, 25.77±0.07°C and 28.08±0.12 μmol photons m^-2^ s^-1^, respectively. Culture containers were placed on a single incubator shelf and randomly rearranged after each clean to minimize confounding by position.

The maximum dark adapted quantum yield (Fv/Fm) of photosystem II (PSII) of *in hospite* Symbiodiniaceae provides an indication of photosynthetic performance and is useful as a bio-monitoring tool (Howe et al. 2017). Fv/Fm is measured as the difference in fluorescence produced by PSII reaction centers when either saturated by intense light (Fm) or in the absence of light (Fo), versus Fm (i.e. (Fm–Fo)/Fm, or Fv/Fm). Symbiodiniaceae Fv/Fm was measured weekly over nine weeks for each culture container after 4 hr into the incubator light cycle and 30 mins dark adaption using imaging pulse amplitude modulated (iPAM) fluorometry (IMAG-MAX/L, Waltz, Germany). Settings for all measurements were: saturating pulse intensity 8, measuring light intensity 2 with frequency 1, actinic light intensity 3, damping 2, gain 2. Average Fv/Fm values for each dish were calculated from readings taken on three to five anemones (Online Resource 3).

Anemones from each culture container were individually homogenized in a sterile glass homogenizer in 1 mL fRSS, and 100 μL of homogenate was collected and stored at −20 °C for total protein measurement. The remaining 900 μL of homogenate was centrifuged at 5000 *g* for 5 min at 4°C to pellet the Symbiodiniaceae while leaving the anemone cells in suspension. A volume of 100 μL of supernatant was collected and stored at –20 °C for host protein measurement. The pelleted Symbiodiniaceae were twice washed with 500 μL fRSS and centrifuged at 5000 *g* for 5 min at 4 °C, and the final pellet resuspended in 500 μL fRSS and stored at –20 °C for Symbiodiniaceae counts. Triplicate cell counts (cells mL^−1^) were completed within 48 hrs of sample collection on a Countess II FL automated cell counter (Life Technologies) and normalized to host protein (mg mL^−1^). This process was repeated 15 times over the course of nine weeks for a total of 45 replicates per genotype.

Samples for protein analysis stored at –20 °C (see above) were used within one month of collection to determine total and host protein (mg mL^−1^) by the Bradford assay (Bradford 1976) against bovine serum albumin standards (Bio-Rad 500-0207). Readings were taken at 595 nm (EnSpire microplate reader MLD2300).

Fv/Fm values and Symbiodiniaceae cell densities were assessed for genotype-specific responses. All variables were tested for normality and homoscedasticity prior to parametric analyses, which were completed in R (v 3.6.0). Genotypic responses of Fv/Fm were compared using a linear model in the R package nlme (Pinheiro et al. 2017) to evaluate independent and interaction relationships between the factors of genotype and time. F-statistics were obtained using the analysis of variance (ANOVA) function, nlme, and pairwise *post hoc* analyses were performed using the glht function in the R package multcomp (Hothorn et al. 2016) with Tukey’s correction for multiple comparisons. Symbiodiniaceae cell densities were pooled across time by genotype and analyzed with a one-way ANOVA with a *post hoc* Tukey test to determine pairwise significance.

## Results and Discussion

### 3.1 Anemones

The assembled 18S rRNA gene sequences of two genotypes (AIMS2 and AIMS4) covered 1591 and 1594 bp, respectively, with two mismatches between the sequences. BLASTn against the NCBI database identified the samples as either *Aiptasia pulchella* or *Exaiptasia pallida. A. pulchella* has been synonymised with *E. pallida* (Grajales and Rodríguez, 2014). However, since *E. diaphana* has precedence (ICZN 2017), all samples were designated *E. diaphana*.

Four SNP genotypes of *E. diaphana*, which were of GBR-origin but AIMS-sourced, were identified and designated AIMS1, AIMS2, AIMS3 and AIMS4 (Fig. 1). The clonal distribution of Euclidean distances was identified (range of 0-9.61) and pairs of individuals with Euclidean distances within this distribution were inferred to have the same genotype (Online Resource 3-4). The Euclidean distance between the originally named Ed.11a and Ed.11b (replicates from the same individual; assigned AIMS3) is in the upper percentiles of the defined clonal distribution, confirming this cut-off is valid.

**Figure 1:**
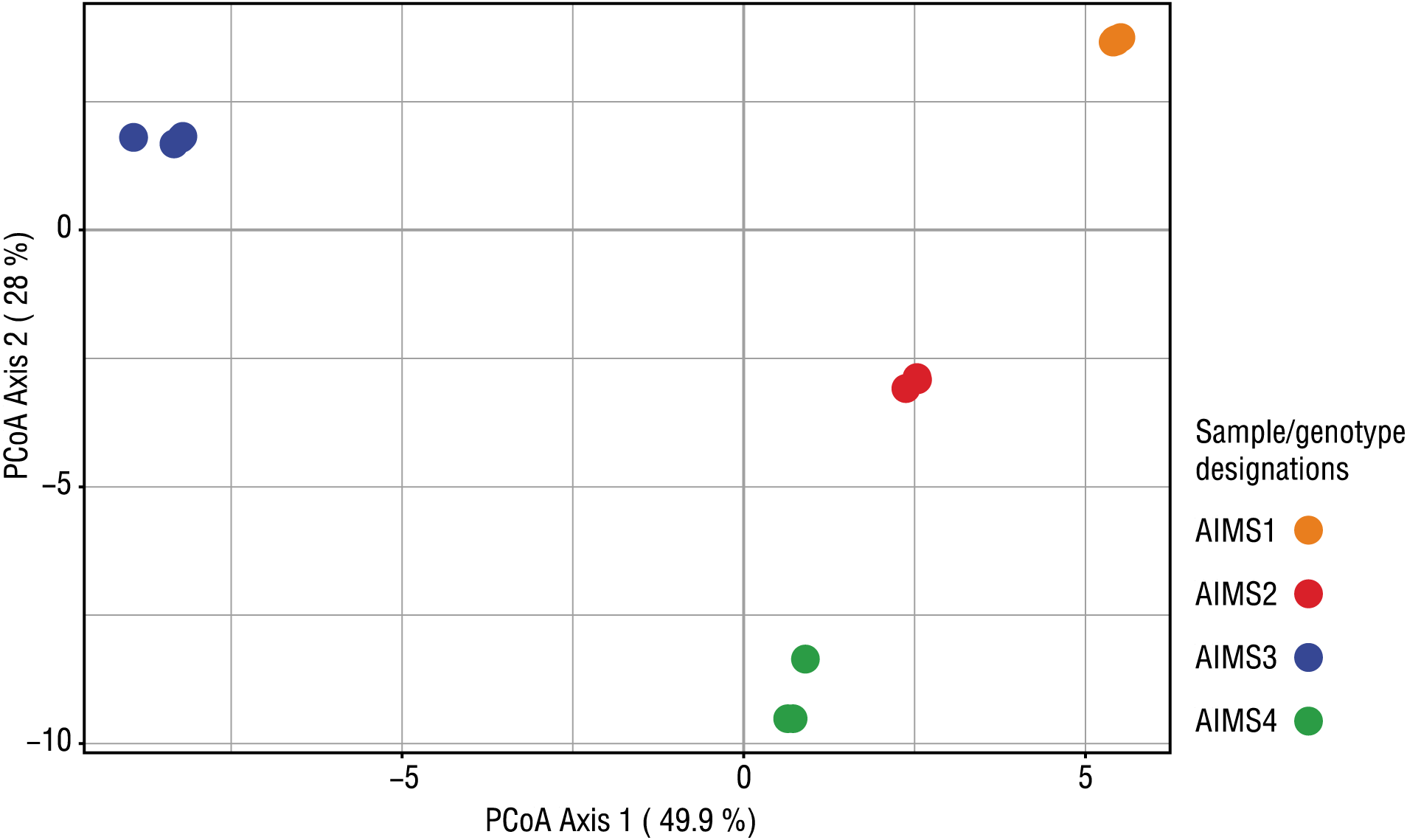
PCoA ordination of *Exaiptasia diaphana* genotypes based on Euclidean genetic distance measurements of SNP data using allele frequencies within individuals to calculate genetic distance between them (AIMS1, n=8; AIMS2, n=5, AIMS3, n=7; AIMS4, n=3). Individuals of the same genotype may overlap in the plot due to their high similarity.

Using phylogenetic analysis with our data, previously described SCAR marker 5, and the *Exaiptasia*-specific gene sequence data (Thornhill et al. 2013; Grawunder et al. 2015; Bellis and Denver 2017), we placed the GBR anemones into a phylogeographical context. The alignment of SCAR5 sequences was 706 bp long and contained 19 variable nucleotide positions (Fig. 2), while the concatenated *Exaiptasia*-specific gene sequences were 3276 bp long with 92 variable nucleotide positions (Fig. 3).

**Figure 2:**
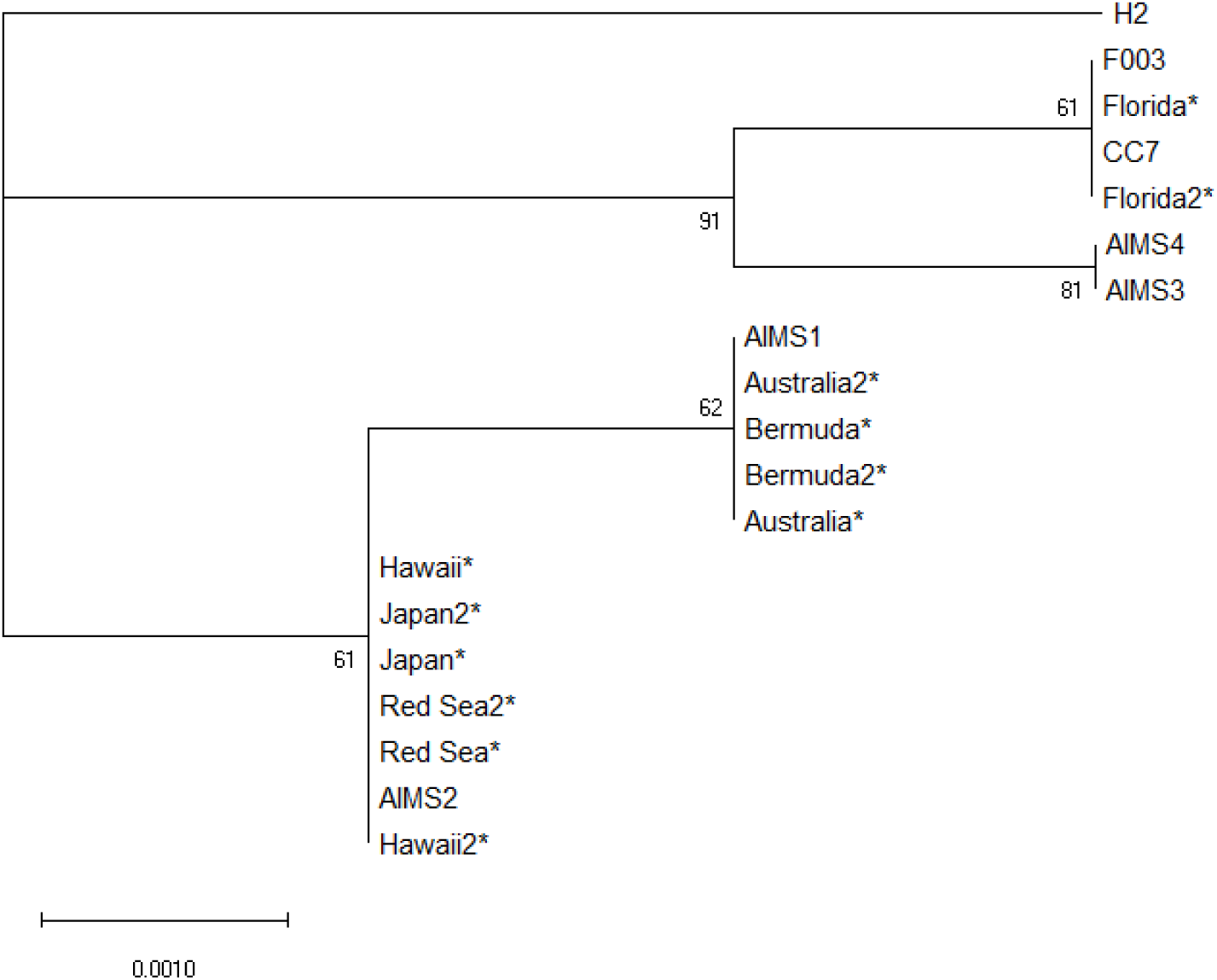
The phylogenetic relationships of the four GBR *E. diaphana* compared to other conspecific anemones sampled across the globe inferred from SCAR marker 5 using the Maximum Likelihood method and General Time Reversible model (Nei and Kumar 2000). The tree with the highest log likelihood is shown with bootstrap values next to the nodes. Initial trees for the heuristic search were obtained automatically by applying the Maximum Parsimony method. The tree is drawn to scale, with branch lengths measured in the number of substitutions per site. This analysis involved 19 *E. diaphana* sequences and 706 nucleotide positions were included. Sequences were from our *E. diaphana* genotypes (AIMS1-4), globally distributed *E. diaphana* from Thornhill et al. (2013), indicated with an asterisk, and from experimental genotypes CC7, H2, and F003 (Grawunder et al. 2015).

**Figure 3:**
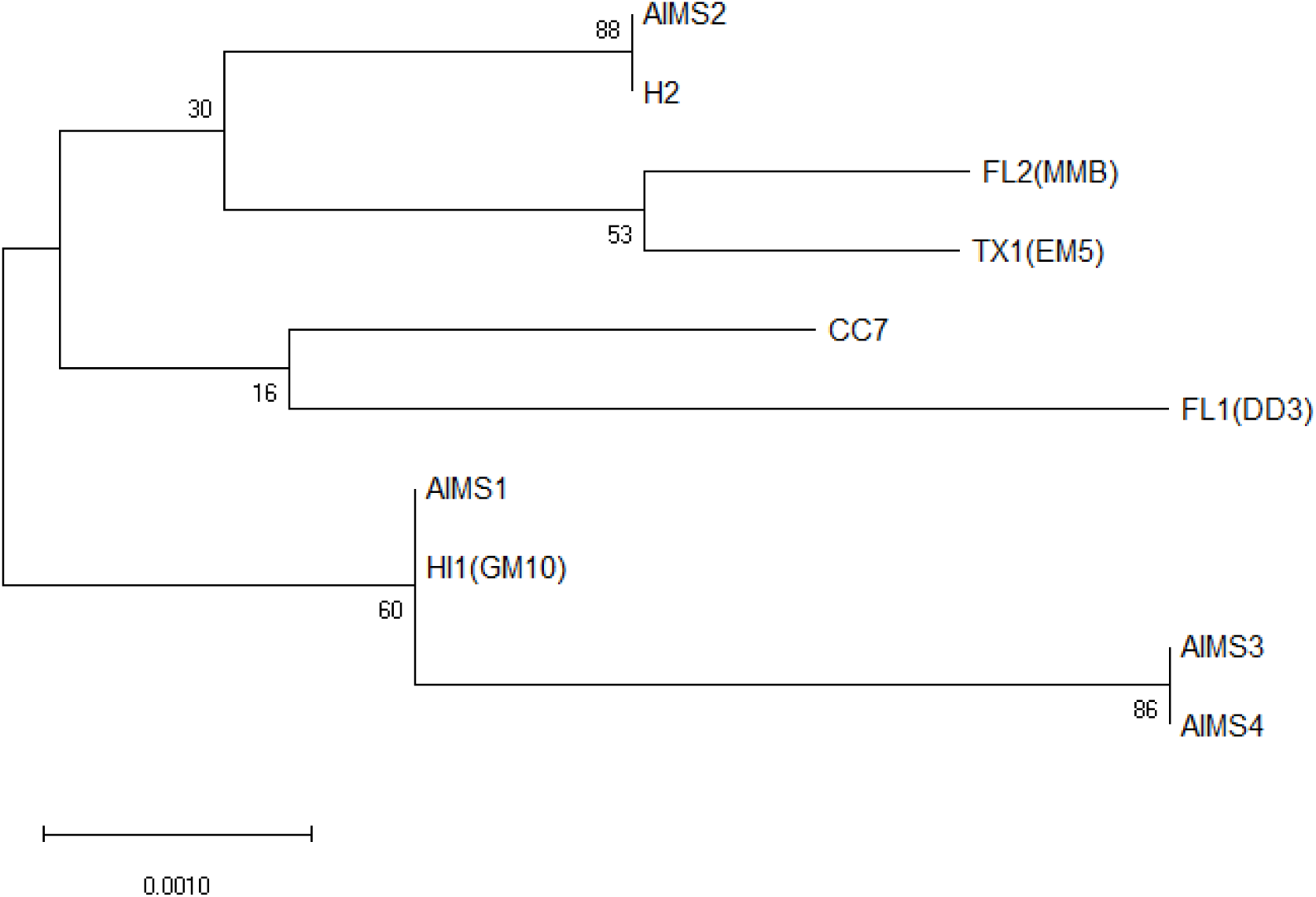
The phylogenetic relationships of the four GBR *E. diaphana* (AIMS1-4) compared to other experimental anemones using the Maximum Likelihood method and General Time Reversible model (Nei and Kumar 2000) on six concatenated *Exaiptasia*-specific gene sequences (Bellis and Denver 2017). Data from Bellis and Denver (2017) was downloaded from GenBank (accession numbers KU847812-KU847847); tree branches are named as strain name followed by the alternative strain name in parentheses. Bootstrap values are shown next to the nodes. Initial trees for the heuristic search were obtained automatically by applying the Maximum Parsimony method. The tree is drawn to scale, with branch lengths measured in the number of substitutions per site. This analysis involved 10 *E. diaphana* sequences and 3267 nucleotide positions were included.

Based on SCAR marker data, *E. diaphana* is regarded as a single pan-global species with two distinct genetic lineages: one from the USA Atlantic coast, and a second consisting of all other *E. diaphana* worldwide (Thornhill et al. 2013). The SCAR5 allele sequenced for genotype AIMS1 is identical with that from anemones originally collected off Heron Island, Australia and Bermuda, while that of AIMS2 is nearly identical to (two base pair differences, both with an ambiguous base) samples from Hawaii, Japan and the Red Sea. AIMS3 and AIMS4 are most closely related to anemones from Florida and North Carolina, USA and are identical to one another in this region.

Similar to Bellis and Denver (2017), our data from the *Exaiptasia*-specific gene sequences (Fig. 3) show that, while anemone strains are genetically distinct, there is not a strong phylogenetic separation between individuals collected from distant geographic locations. Again, AIMS1-4 show genetic variation, with AIMS1 and AIMS2 clustering with anemone strains originating from Coconut Island, Hawaii, USA which were collected independently in 1979 and early 2000’s, respectively. Corresponding with the SCAR5 loci data, AIMS3-4 have near identical sequences in the sequenced regions (differences at two heterozygous sites). Because the available data for SCAR5 and the *Exaiptasia*-specific gene targets are not from all the same individuals, with the exception of CC7 and H2, comparing between the two is not feasible. However, given the larger number of alignment positions and variable sites in the gene regions with better PCR results, we suggest that researchers use the primers presented in Bellis and Denver (2017) for future comparisons between *E. diaphana* used in experiments. The diversity that is revealed by whole genome SNP analysis (Fig. 1) is hidden with these six *Exaiptasia*-specific gene sequences, suggesting that there are more informative loci not yet published for *E. diaphana* genotyping.

There are several possible explanations for the AIMS1-4 anemones to be spread throughout the phylogenetic trees. First, due to small sample sizes at all sampling locations, only a subset of the alleles have been sampled and location-specific alleles may have been missed. Second, it is conceivable that the GBR is the source of all other *E. diaphana* populations and founder effects mean that other geographic locations have *E. diaphana* that represent only some of the diversity. Third, it is possible that the GBR *E. diaphana* was a distinct lineage and the GBR has since been invaded by *E. diaphana* from other lineages, or the GBR lineage has been introduced elsewhere. Introductions over such vast spatial scales may have occurred via ship ballast water or attached to ships hulls in fouling biomass, which is notorious for transporting marine life and introducing invasive animals and plants, or via the aquarium trade or marine farms. Irrespective of the cause, the genetic variation across the four GBR genotypes is more representative of global diversity than a single localized population. Because strain-specific responses to environmental variables have been observed among *E. diaphana* strains (Bellis and Denver 2017; Cziesielski et al. 2018) it is critical to conduct experiments, such as those regarding climate change and symbiosis, with a diverse set of individuals, much like the diversity presented by the GBR-sourced *E. diaphana*.

### 3.2 Symbiosis with Symbiodiniaceae

According to a global survey by Thornhill et al. (2013), Pacific Ocean *E. diaphana* (e.g., H2) associate exclusively with the Symbiodiniaceae species *Breviolum minutum* (formerly ITS2 type *Symbiodinium* Clade B, sub-clade B1 (LaJeunesse et al. 2018)). Symbiodiniaceae sequences from the four GBR *E. diaphana* genotypes were almost exclusively *Breviolum minutum* (>99.6%), thus concurring with Thornhill et al. (2013). Two as-yet unnamed species of *Breviolum* (previously known as *Symbiodinium* sub-clades B1i and B1L (LaJeunesse et al. 2018)) were also identified in *E. diaphana*. Their very low relative abundance suggests they are either intragenomic variants or rare strains.

Stable endosymbiotic relationships between corals and Symbiodiniaceae are vital for sustaining coral reef ecosystems; this symbiotic relationship is the focus of much coral reef research. Interestingly, while the GBR anemones are genetically diverse, all host *B. minutum* as their sole Symbiodiniaceae. Given this, we may be able to investigate the symbiotic mechanisms of the host-algal relationship and answer questions about this symbiosis that are not restricted by anemone strain.

### 3.3 Sex of *E. diaphana*

*E. diaphana* lacks obvious gender defining morphological features, but gonad development is related to size (Chen et al. 2008). Partially developed oocytes with germinal vesicles (*i.e.*, the nucleus of an oocyte arrested in prophase of meiosis I) were observed in histological slides in animals of AIMS1, AIMS3 and AIMS4 and Stage V spermaries were observed in AIMS2 (Fig. 4). In Stage V, the spermary is made up of a mass of spermatozoa with their tails facing in the same direction. In this advanced stage the spermatozoa are capable of fertilization (Fadlallah and Pearse 1982; Goffredo et al. 2012). While most AIMS1 anemones grew to only 5 mm in pedal disk diameter, gonad development was still observed. Differences between male and female gonads were present but were macroscopically cryptic. Male gonads have been observed to be smaller and lighter colored than female gonads in *E. diaphana* (Grawunder et al. 2015), and this was also the case in the GBR-sourced females (AIMS1, AIMS3 and AIMS4) and male (AIMS2) *E. diaphana*.

**Figure 4:**
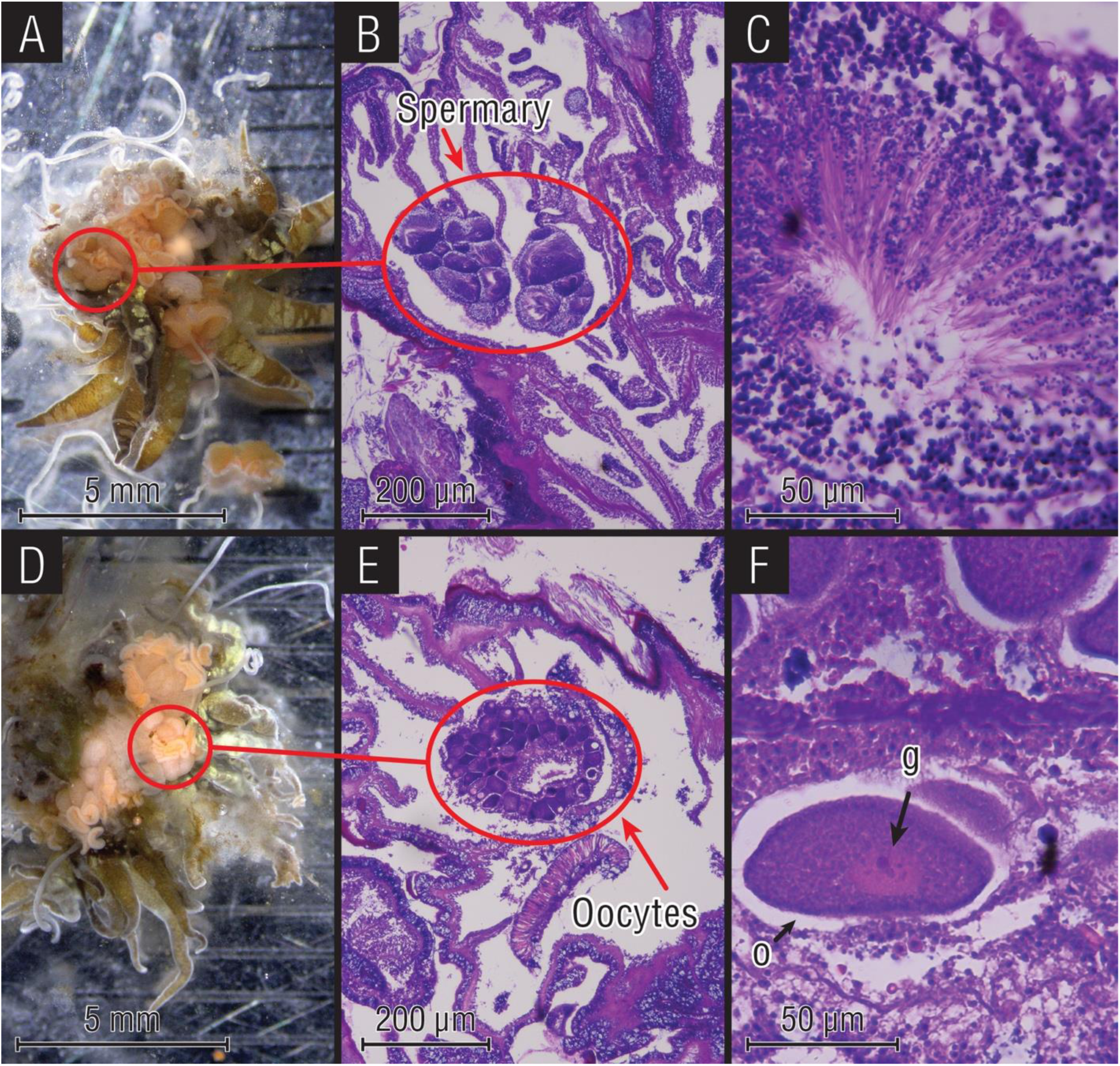
(A) Dissected male anemone with developing gonads (AIMS2, ∼1 cm pedal disk diameter); (B) H&E stained tissue section, with stage 5 spermary; (C) Increased magnification of stage 5 spermary; (D) dissected female anemone with developed gonads (AIMS4, ∼1 cm pedal disk diameter); (E) H&E stained tissue section with oocytes; (F) increased magnification of female gonad with developing oocyte (o) and germinal vesicle (g).

### 3.4 *E. diaphana* and Symbiodiniacease Physiology

Throughout the nine-week evaluation period all *E. diaphana* maintained a healthy appearance, including *in hospite* Symbiodiniaceae, tentacle extension, active feeding of *A. salina* nauplii and asexual propagation. Average Symbiodiniaceae cell densities (normalized to host protein) were significantly different between genotypes (one-way ANOVA (F(3,200)=3.985, p=0.00872; Fig. 5; Online Resource 6). All anemones hosted ∼10^6^ Symbiodiniaceae cells mg^-1^ host protein, which is comparable to densities reported for other lab cultured model *E. diaphana* systems (Hoadley et al. 2015; Hawkins et al. 2016b; Rädecker et al. 2018) and is similar to Symbiodiniaceae cell densities found in scleractinian corals (Cunning and Baker 2014; Ziegler et al. 2015; Kenkel and Bay 2018). The only statistically significant difference observed was the lower Symbiodiniaceae cell density of AIMS2 (mean ± SE; 7.17 x 10^6^ ± 2.73 x 10^5^ per mg host protein, n=66) compared to AIMS3 (mean ± SE; 8.46 x 10^6^ ± 2.96 x 10^5^ per mg host protein, n=66) (Fig. 5).

**Figure 5:**
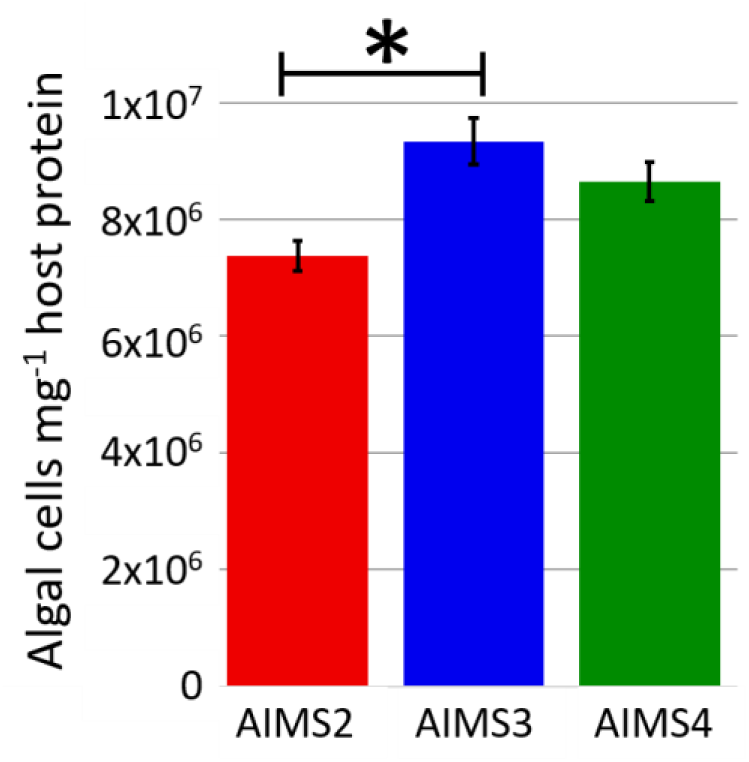
Mean ± 1SEM Symbiodiniaceae cells mg^−1^ host protein for GBR *E. diaphana* genotypes AIMS2-4. Seventy-five AIMS2-4 anemones were collected over a period of nine weeks. Asterisk indicates significant difference, *p*<0.05.

All genotypes experienced a nearly identical drop in Fv/Fm after the light intensity was increased from 12 to 28 μmol photons m^-2^ s^-1^, from an average of 0.53 on day 0, to an average of 0.40 by day 21 (Online Resource 7; Fig. 6). However, by day 36 the Fv/Fm values had returned to initial levels.

**Figure 6:**
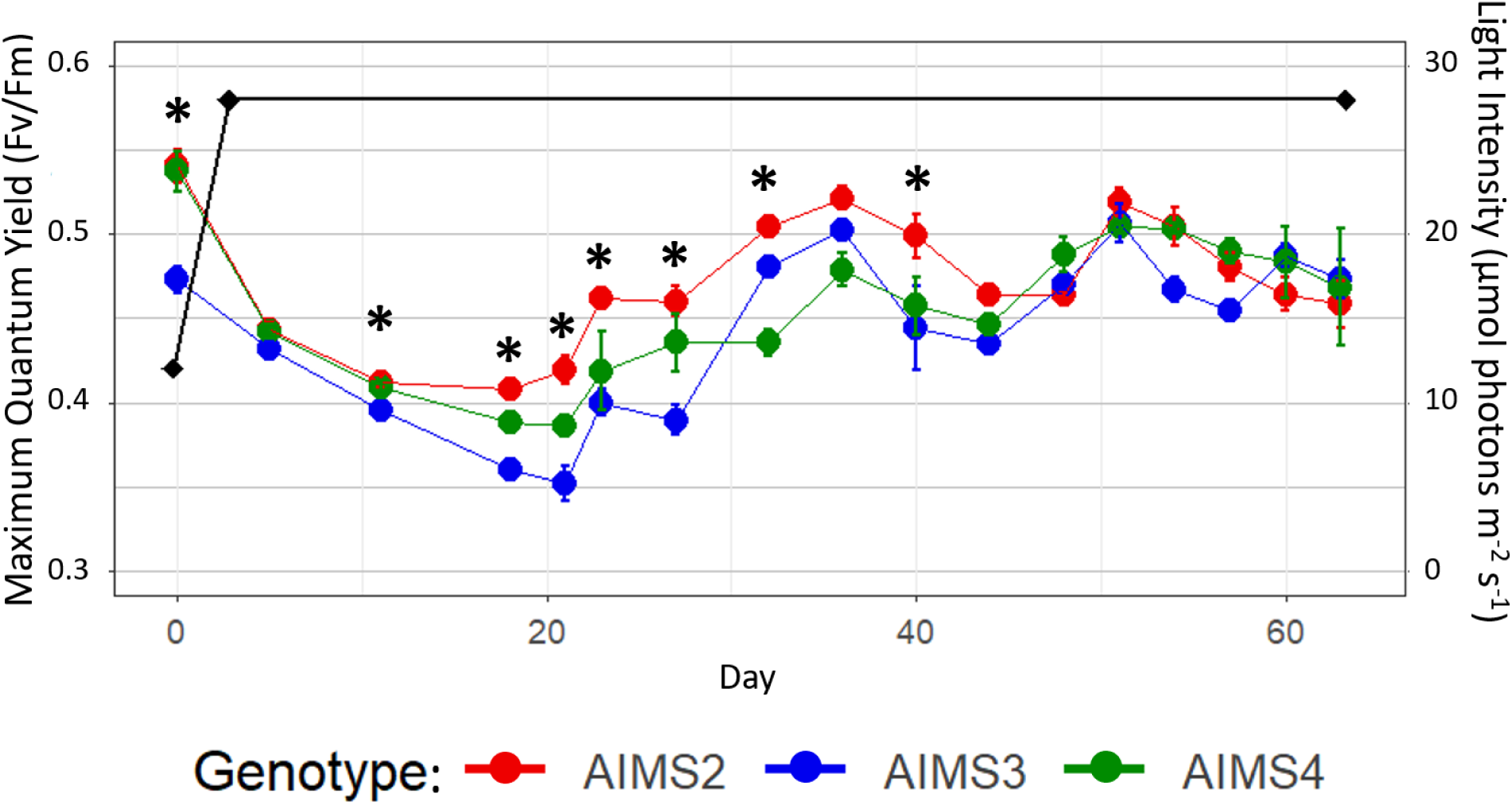
Fv/Fm measurements for anemones AIMS2, AIMS3 and AIMS4 over a 63-day period. Anemones were initially exposed to light levels (black line) of 12 μmol photons m^-2^ s^-1^ (12:12 light:dark cycle), which were then increased to 28 μmol photons m^-2^ s^-1^ over a 72 h period. Day 0 marks the start of light ramping. The anemones took ∼35 d to recover their maximum quantum yield due to the increase in light exposure although Symbiodiniaceae densities remained largely constant. Asterisks indicate significant differences in pairwise comparisons between genotypes at given time points (Online Resource 8)

Changes in environmental variables, such as light intensity, are known to influence photobiological behavior of Symbiodiniaceae (Wangpraseurt et al. 2014; Hoadley and Warner 2017). A change in light intensitycan alter photosynthetic efficiency as measured by Fv/Fm (Hoadley and Warner 2017). GBR anemone genotypes AIMS2, AIMS3 and AIMS4 took 36 d to recover from this environmental change and return to photosynthetic efficiencies recorded at 12 μmol photons m^-2^ s^-1^. Such information is critical for planning and conducting symbiosis studies. Most of the statistical differences between the genotypes occurred between days 19-28 (Online Resource 8) when Fv/Fm was lowest. Once acclimated at day 36, average Fv/Fm values for all genotypes were not significantly different (mean ± SE; 0.479 ± 0.003, n=80), except for day 40 where the Fv/Fm of AIMS2 was significantly higher than AIMS3 (*p*=0.0068).

As all the GBR-sourced anemones harbour *Breviolum minutum* as their homologous symbiont type, it is not unexpected that the different genotypes would have similar maximum Fv/Fm values. Furthermore, similar Fv/Fm values have been reported for other anemones hosting homologous *B. minutum* (Hawkins et al. 2016b; Hillyer et al. 2017). However, it is noteworthy that AIMS2 is not only able to recover its photosynthetic efficiency quicker from changing light levels with a milder dip in Fv/Fm values (Fig. 6), but also hosts significantly fewer Symbiodiniaceae cells mg^-1^ host protein compared to AIMS3 (Fig. 5). Reductions in the maximum quantum yield of photosystem II (PSII) (Fv/Fm) is observed in the early phases of natural bleaching events (Gates et al. 1992; Franklin et al. 2004) and the ability of AIMS2 to maintain a higher efficiency of PSII photochemistry during changing environmental conditions could translate into higher thermal tolerance (Suggett et al. 2008; Ragni et al. 2010; Goyen et al. 2017).

These individuals are genetically diverse based on SNP genotyping (Fig. 1) and phylogenetic analysis (Fig. 2-3); the phenotypic feature of AIMS2 being more robust to increasing light levels compared to AIMS3 and AIMS4 could be a host genotypic effect. There is evidence that genetic variation of *E. diaphana* may influence holobiont response to heat stress, though this hypothesis has only been tested on anemone strains hosting different Symbiodiniaceae species (Bellis and Denver 2017; Cziesielski et al. 2018) or after experimentally bleaching anemones and inoculating with new heterologous algal cells (Perez et al. 2001). As we have four GBR-sourced *E. diaphana* genotypes with inherent genetic variability and all contain *B. minutum* as their homologous symbiont, we will be able to explore the roles of host and symbiont in the bleaching response.

An alternative possibility is that the GBR anemones’ Symbiodiniaceae communities comprise diversity that may be hidden under the resolution of the ITS2 sequences we used in this experiment, which are driving the differences in photosynthetic efficiency. Distinct strains of a given Symbiodiniaceae species can have different susceptibilities to thermal stress (Ragni et al. 2010; Howells et al. 2012; Hawkins et al. 2016a), with evidence that these variations in thermal optima can drive host-Symbiodiniaceae interactions (Hawkins et al. 2016a). Thus, varying rates of recovery of Fv/Fm among Symbiodiniaceae strains (Fig. 6) could provide a mechanism for the emergence of novel and potentially resilient cnidarian-Symbiodiniaceae associations in a rapidly warming environment.

Another explanation for the physiological differences between AIMS2 and AIMS3 (Fig. 5) is through algal cell density moderation by the host. It is thought that the coral host controls Symbiodiniaceae densities through nitrogen limitation (Falkowski et al. 1993), although the mechanisms are not well understood (Davy et al. 2012). During temperature stress, higher densities of Symbiodiniaceae have been implicated in increasing the susceptibility of corals to bleaching, potentially as a result of the higher reactive oxygen species production relative to corals’ antioxidant capacity (Cunning and Baker 2012). Altogether, our data suggest that AIMS2, which hosted fewer algal symbionts and recovered from increased light conditions faster than AIMS3, may be more resilient to thermal stress, while AIMS3 could be more susceptible to bleaching with AIMS4 as an intermediate.

## 4. Conclusions and Future Directions

The study of *E. diaphana* anemones of GBR origin described here provide further information on phenotypic and genetic variation within this species and complements data on the more widely used *E. diaphana* strains CC7 and H2. The four genotypes in our collections capture a level of genetic diversity previously observed in animals from different oceans and are therefore a hugely valuable addition to the model collections. Knowledge of their characteristics enhances and broadens the potential of this model system for climate change research in corals, particularly, but not exclusively, for Australian researchers. We propose future research on this collection should focus on characterization of associated prokaryotes to explore the value of these animals as models for coral-prokaryote symbiotic interactions. Future research in cnidarian-prokaryotic interactions would be enhanced by the development of axenic (germ-free) or gnotobiotic (with a known microbial community) *E. diaphana* cultures. They could be used to test the influence of native and non-native microbiota on holobiont performance, and the ability of probiotic inocula to support animal health during stress (Alagely et al. 2011; Damjanovic et al. 2017; Rosado et al. 2018).

## Supporting information

Online Resource 4

Online Resource 3

Online Resource 2

Online Resource 8

Online Resource 7

Online Resource 6

Online Resource 1

Online Resource 5

## 5. Acknowledgments

This research was funded by Australian Research Council Discovery Project grants DP160101468 (to MJHvO and LLB) and DP160101539 (to GIM and MJHvO). We thank Lesa Peplow for facilitating transport of the initial anemone cultures from AIMS to SUT and UoM and Rebecca Alfred from SUT for initial anemone culture maintenance. We acknowledge Anton Cozijnsen, Keren Maor-Landaw, Samantha Girvan, Ruby Vanstone and Gabriela Rodriguez from University of Melbourne for assisting with anemone husbandry and Laura Leone, Lisa Foster, and Lona Dinha from the Melbourne Histology Platform for histological sample preparation and sectioning. SCAR marker reference sequences were provided by Dan Thornhill and Liz Hambleton (AG Guse Lab, Centre for Organismal Studies (COS), Universität Heidelberg). MJHvO acknowledges Australian Research Council Laureate Fellowship FL180100036.

