## Supplementary material for "*Exaiptasia diaphana* from the Great Barrier Reef: a valuable resource for coral symbiosis research": Online Resource 4

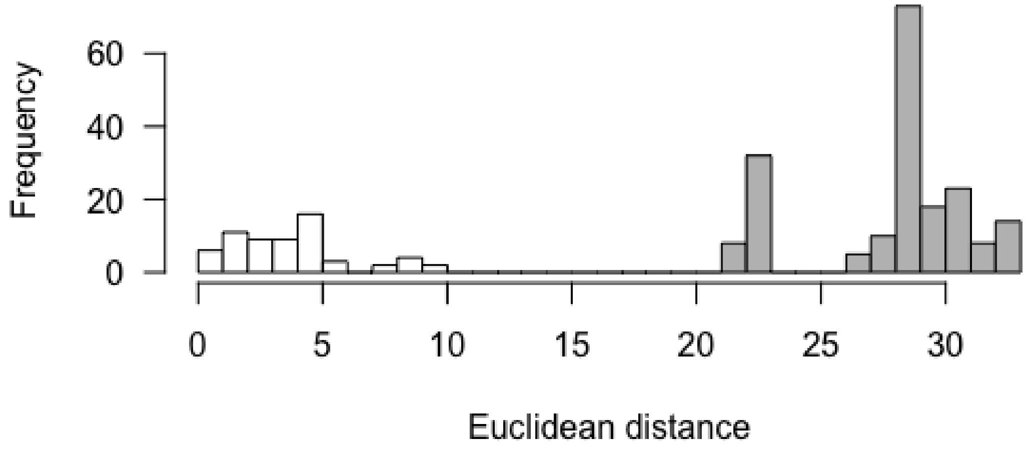


**Online Resource 4:** Histogram of Euclidean distances among *Exaiptasia diaphana* individuals. The inferred clonal (range of 0-9.61) and inter-genotypic distributions are shown in white and grey, respectively.
