## Supplementary material for "*Exaiptasia diaphana* from the Great Barrier Reef: a valuable resource for coral symbiosis research": Online Resource 3

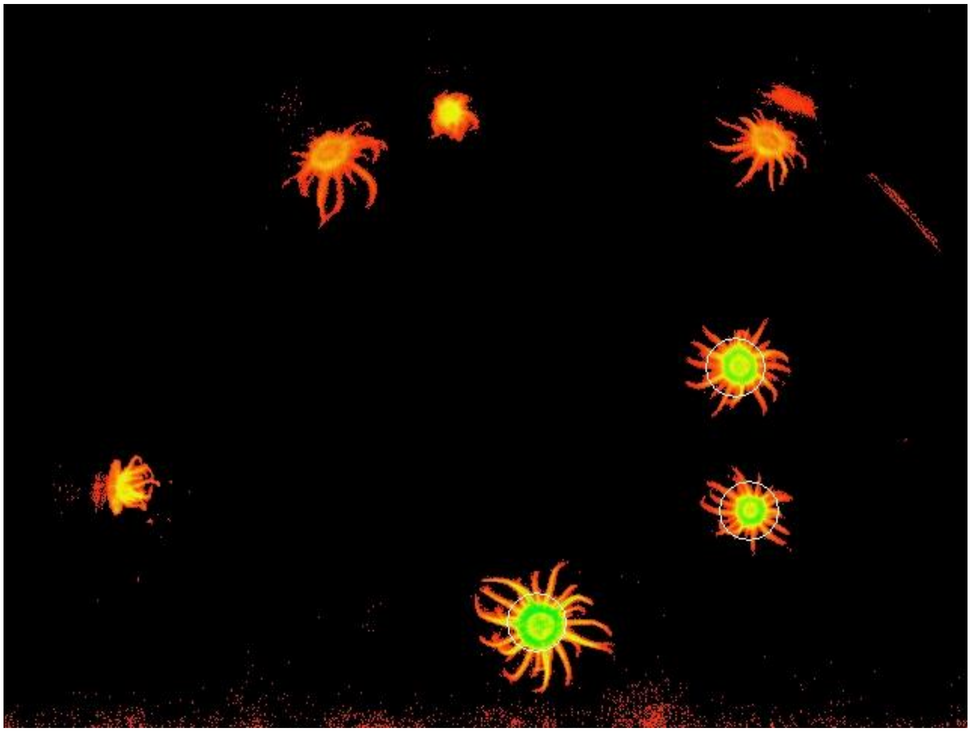

**Online Resource 3:** Areas of interest (AOIs) are show as white circles surrounding the central body of the anemones and including the proximal portion of the tentacles. A minimum of three anemones per jar were selected, avoiding individuals on the sides that were out of focus and could skew the data. $F_0$ values ranged from 0.100 to 0.300. No scale available.
