## Supplementary material for "*Exaiptasia diaphana* from the Great Barrier Reef: a valuable resource for coral symbiosis research": Online Resource 2

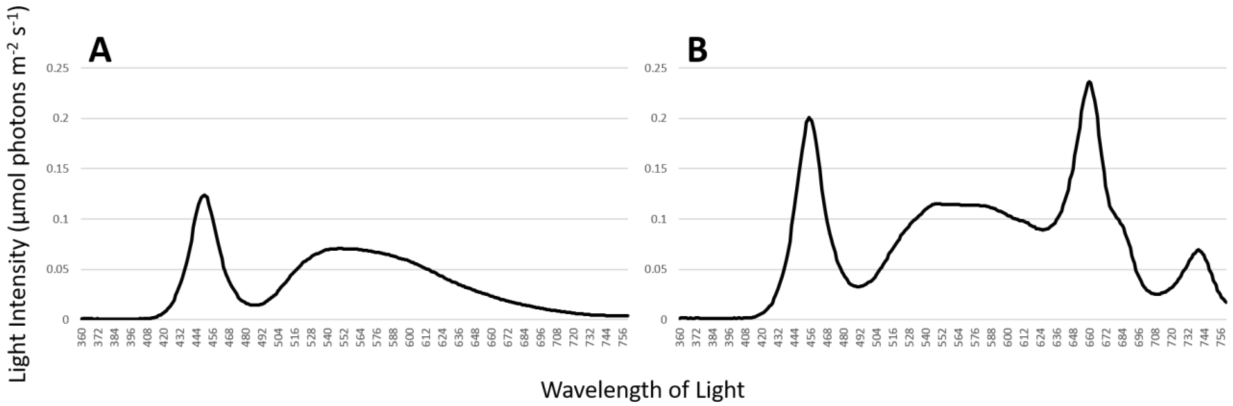


**Online Resource 2:** Spectra of light intensities (HiPoint HR-350 LED meter) from the A) walk-in incubator where the GBR-sourced stock anemones are held (fitted with white LEDs only) and B) the experimental growth chambers (Taiwan HiPoint Corporation model 740FHC) fitted with red, white, and infrared LED lights.
