## Supplementary material for "*Exaiptasia diaphana* from the Great Barrier Reef: a valuable resource for coral symbiosis research": Online Resource 8

**Online Resource 8:** Pairwise comparisons between genotypes by sampling day from a linear mixed effects model with genotype and sampling day as fixed effects and anemone jar as a random effect. **Significant differences are noted in bold.**

| Sampling Day | Genotype | Mean | SE | lower.CL | upper.CL | contrast | estimate | SE | df | t.ratio | p.value |
|---|---|---|---|---|---|---|---|---|---|---|---|
| 0 | 2 | 0.541 | 0.01202 | 0.517 | 0.565 | 2-3 | 0.067667 | 0.01701 | 60 | 3.979 | **0.0005** |
| 0 | 3 | 0.473 | 0.01202 | 0.449 | 0.497 | 2-4 | 0.003333 | 0.01701 | 60 | 0.196 | 0.979 |
| 0 | 4 | 0.537 | 0.01202 | 0.513 | 0.561 | 3-4 | -0.06433 | 0.01701 | 60 | -3.783 | **0.001** |
| 5 | 2 | 0.441 | 0.00721 | 0.427 | 0.456 | 2-3 | 0.006964 | 0.00992 | 60 | 0.702 | 0.7634 |
| 5 | 3 | 0.434 | 0.00682 | 0.421 | 0.448 | 2-4 | -0.00335 | 0.00992 | 60 | -0.338 | 0.9391 |
| 5 | 4 | 0.445 | 0.00682 | 0.431 | 0.458 | 3-4 | -0.01032 | 0.00964 | 60 | -1.07 | 0.5361 |
| 11 | 2 | 0.412 | 0.00491 | 0.402 | 0.422 | 2-3 | 0.016889 | 0.00694 | 60 | 2.433 | **0.0466** |
| 11 | 3 | 0.395 | 0.00491 | 0.385 | 0.405 | 2-4 | 0.003278 | 0.00694 | 60 | 0.472 | 0.8846 |
| 11 | 4 | 0.409 | 0.00491 | 0.399 | 0.419 | 3-4 | -0.01361 | 0.00694 | 60 | -1.961 | 0.131 |
| 18 | 2 | 0.408 | 0.00491 | 0.398 | 0.418 | 2-3 | 0.0477 | 0.00694 | 60 | 6.871 | **<.0001** |
| 18 | 3 | 0.36 | 0.00491 | 0.351 | 0.37 | 2-4 | 0.020478 | 0.00694 | 60 | 2.95 | **0.0124** |
| 18 | 4 | 0.388 | 0.00491 | 0.378 | 0.398 | 3-4 | -0.02722 | 0.00694 | 60 | -3.921 | **0.0007** |
| 21 | 2 | 0.421 | 0.00831 | 0.404 | 0.437 | 2-3 | 0.068978 | 0.01175 | 60 | 5.869 | **<.0001** |
| 21 | 3 | 0.352 | 0.00831 | 0.335 | 0.368 | 2-4 | 0.034306 | 0.01175 | 60 | 2.919 | **0.0135** |
| 21 | 4 | 0.386 | 0.00831 | 0.37 | 0.403 | 3-4 | -0.03467 | 0.01175 | 60 | -2.95 | **0.0124** |
| 23 | 2 | 0.463 | 0.00831 | 0.447 | 0.48 | 2-3 | 0.063478 | 0.01175 | 60 | 5.401 | **<.0001** |
| 23 | 3 | 0.4 | 0.00831 | 0.383 | 0.417 | 2-4 | 0.044639 | 0.01175 | 60 | 3.798 | **0.001** |
| 23 | 4 | 0.419 | 0.00831 | 0.402 | 0.435 | 3-4 | -0.01884 | 0.01175 | 60 | -1.603 | 0.2523 |
| 27 | 2 | 0.461 | 0.00831 | 0.445 | 0.478 | 2-3 | 0.071811 | 0.01175 | 60 | 6.11 | **<.0001** |
| 27 | 3 | 0.39 | 0.00831 | 0.373 | 0.406 | 2-4 | 0.025473 | 0.01175 | 60 | 2.167 | 0.0852 |
| 27 | 4 | 0.436 | 0.00831 | 0.419 | 0.453 | 3-4 | -0.04634 | 0.01175 | 60 | -3.943 | **0.0006** |
| 32 | 2 | 0.508 | 0.01155 | 0.485 | 0.531 | 2-3 | 0.021037 | 0.01634 | 60 | 1.288 | 0.4075 |
| 32 | 3 | 0.487 | 0.01155 | 0.463 | 0.51 | 2-4 | 0.06324 | 0.01634 | 60 | 3.871 | **0.0008** |
| 32 | 4 | 0.444 | 0.01155 | 0.421 | 0.467 | 3-4 | 0.042203 | 0.01633 | 60 | 2.584 | **0.0323** |
| 36 | 2 | 0.523 | 0.01398 | 0.495 | 0.551 | 2-3 | 0.014644 | 0.01813 | 60 | 0.808 | 0.6998 |
| 36 | 3 | 0.509 | 0.01155 | 0.485 | 0.532 | 2-4 | 0.035513 | 0.01813 | 60 | 1.959 | 0.1314 |
| 36 | 4 | 0.488 | 0.01155 | 0.465 | 0.511 | 3-4 | 0.02087 | 0.01633 | 60 | 1.278 | 0.4132 |

| | | | | | | | | | | | |
|---|---|---|---|---|---|---|---|---|---|---|---|
| 40 | 2 | 0.502 | 0.01155 | 0.479 | 0.525 | 2-3 | 0.051704 | 0.01634 | 60 | 3.165 | **0.0068** |
| 40 | 3 | 0.45 | 0.01155 | 0.427 | 0.473 | 2-4 | 0.035907 | 0.01634 | 60 | 2.198 | 0.0797 |
| 40 | 4 | 0.466 | 0.01155 | 0.443 | 0.489 | 3-4 | -0.0158 | 0.01633 | 60 | -0.967 | 0.6003 |
| 44 | 2 | 0.467 | 0.01155 | 0.444 | 0.49 | 2-3 | 0.025704 | 0.01634 | 60 | 1.574 | 0.2649 |
| 44 | 3 | 0.441 | 0.01155 | 0.418 | 0.464 | 2-4 | 0.01124 | 0.01634 | 60 | 0.688 | 0.7713 |
| 44 | 4 | 0.455 | 0.01155 | 0.432 | 0.478 | 3-4 | -0.01446 | 0.01633 | 60 | -0.886 | 0.6515 |
| 48 | 2 | 0.467 | 0.01155 | 0.444 | 0.49 | 2-3 | -0.0093 | 0.01634 | 60 | -0.569 | 0.837 |
| 48 | 3 | 0.476 | 0.01155 | 0.453 | 0.499 | 2-4 | -0.03009 | 0.01634 | 60 | -1.842 | 0.1648 |
| 48 | 4 | 0.497 | 0.01155 | 0.474 | 0.52 | 3-4 | -0.0208 | 0.01633 | 60 | -1.273 | 0.4157 |
| 51 | 2 | 0.522 | 0.01155 | 0.499 | 0.545 | 2-3 | 0.009371 | 0.01634 | 60 | 0.574 | 0.8346 |
| 51 | 3 | 0.513 | 0.01155 | 0.489 | 0.536 | 2-4 | 0.008907 | 0.01634 | 60 | 0.545 | 0.8493 |
| 51 | 4 | 0.513 | 0.01155 | 0.49 | 0.536 | 3-4 | -0.00046 | 0.01633 | 60 | -0.028 | 0.9996 |
| 54 | 2 | 0.508 | 0.01155 | 0.485 | 0.531 | 2-3 | 0.034371 | 0.01634 | 60 | 2.104 | 0.0975 |
| 54 | 3 | 0.473 | 0.01155 | 0.45 | 0.496 | 2-4 | -0.00443 | 0.01634 | 60 | -0.271 | 0.9604 |
| 54 | 4 | 0.512 | 0.01155 | 0.489 | 0.535 | 3-4 | -0.0388 | 0.01633 | 60 | -2.375 | 0.0534 |
| 57 | 2 | 0.484 | 0.01155 | 0.461 | 0.507 | 2-3 | 0.022704 | 0.01634 | 60 | 1.39 | 0.3526 |
| 57 | 3 | 0.461 | 0.01155 | 0.438 | 0.484 | 2-4 | -0.01509 | 0.01634 | 60 | -0.924 | 0.6274 |
| 57 | 4 | 0.499 | 0.01155 | 0.476 | 0.522 | 3-4 | -0.0378 | 0.01633 | 60 | -2.314 | 0.0615 |
| 60 | 2 | 0.467 | 0.01155 | 0.444 | 0.49 | 2-3 | -0.02596 | 0.01634 | 60 | -1.589 | 0.2581 |
| 60 | 3 | 0.493 | 0.01155 | 0.47 | 0.516 | 2-4 | -0.02476 | 0.01634 | 60 | -1.516 | 0.2909 |
| 60 | 4 | 0.492 | 0.01155 | 0.469 | 0.515 | 3-4 | 0.001203 | 0.01633 | 60 | 0.074 | 0.997 |
| 63 | 2 | 0.461 | 0.01155 | 0.438 | 0.484 | 2-3 | -0.01796 | 0.01634 | 60 | -1.1 | 0.5181 |
| 63 | 3 | 0.479 | 0.01155 | 0.456 | 0.502 | 2-4 | -0.01609 | 0.01634 | 60 | -0.985 | 0.589 |
| 63 | 4 | 0.477 | 0.01155 | 0.454 | 0.5 | 3-4 | 0.00187 | 0.01633 | 60 | 0.114 | 0.9928 |
