## Supplementary material for "*Exaiptasia diaphana* from the Great Barrier Reef: a valuable resource for coral symbiosis research": Online Resource 7

**Online Resource 7:** Statistical output from linear mixed effect output where genotype and sampling day were chosen as fixed effects and anemone jar was identified as a random effect.

| | numDF | denDF | F-value | p-value |
|---|---|---|---|---|
| (Intercept) | 1 | 175 | 59220.65 | <.0001 |
| Genotype | 2 | 60 | 31.76 | <.0001 |
| Sampling_Day | 16 | 175 | 70.76 | <.0001 |
| Genotype:Sampling_Day | 32 | 175 | 3.96 | <.0001 |

4
