## Supplementary material for "*Exaiptasia diaphana* from the Great Barrier Reef: a valuable resource for coral symbiosis research": Online Resource 6

**Online Resource 6:** TukeyHSD pairwise comparisons for Symbiodiniaceae cell densities normalized to host protein between genotypes from a one-way ANOVA ($F_{(3,200)}$=3.985, p=0.00872). **Significant differences are noted in bold.**

| *Genotype* | *diff* | *lwr* | *upr* | *p* |
|---|---|---|---|---|
| *2-1* | -2196232 | -4966379 | 573915.5 | 0.172015 |
| *3-1* | -902961 | -3673108 | 1867186 | 0.83311 |
| *4-1* | -1109622 | -3879769 | 1660525 | 0.727473 |
| ***3-2*** | **1293271** | **162362.8** | **2424179** | **0.01784** |
| *4-2* | 1086609 | -44298.5 | 2217517 | 0.064653 |
| *4-3* | -206661 | -1337569 | 924246.6 | 0.964843 |
