## Supplementary material for "*Exaiptasia diaphana* from the Great Barrier Reef: a valuable resource for coral symbiosis research": Online Resource 1

**Online Resource 1:** Histological processing steps used to serially dehydrate anemone tissue after fixation but prior to embedding. This protocol is modified from Carlisle et al. (2017).

| Process | Solution | Duration |
|---|---|---|
| Rinsing | Deionized water | 15 min |
| | Running tap water | 4 hr |
| Dehydration | 50% ethanol | 50 min |
| | 70% ethanol | Overnight |
| | 90% ethanol | 50 min |
| | 100% ethanol I | 50 min |
| | 100% ethanol II | 50 min |
| | 100% ethanol III | 50 min |
| Clearing | 1:1 100% ethanol:Xylene | 50 min |
| | Xylene I | 50 min |
| | Xylene II | 50 min |
| Embedding | Histoplast paraffin wax I | 1 hr |
| | Histoplast paraffin wax II | 1 hr |
| | Histoplast paraffin wax III | 1 hr |
| | Histoplast paraffin wax IV | 1 hr |

Carlisle JF, Murphy GK, Roark AM (2017) Body size and symbiotic status influence gonad development in Aiptasia pallida anemones Symbiosis 71:121-127 doi:10.1007/s13199-016-0456-1
