## Supplementary material for "*Exaiptasia diaphana* from the Great Barrier Reef: a valuable resource for coral symbiosis research": Online Resource 5

**Online Resource 5:** Euclidean distance matrix between *E. diaphana* individuals. Individuals are arranged by genotype (AIMS1-4) and distances within and outside the clonal distribution are shown in white and grey, respectively. Name codes: Ed = *Exaiptasia diaphana* sent from AIMS, S= *Exaiptasia diaphana* originally from AIMS but held at Swinburne University of Technology (with I, II, A, B, and C as individuals)

| **Genotype** | ***AIMS1*** | | | | | | | | ***AIMS3*** | | | | | | | ***AIMS2*** | | | | | ***AIMS4*** | | |
|---|---|---|---|---|---|---|---|---|---|---|---|---|---|---|---|---|---|---|---|---|---|---|---|
| **Individual** | Ed.01 | Ed.02 | Ed.03 | Ed.07 | Ed.08 | Ed.09 | Ed.12 | Ed.21 | Ed.06 | Ed.11a | Ed.11b | Ed.14 | Ed.17 | Ed.18 | Ed.19 | Ed.10 | Ed.20 | Ed.22 | SC | SII | SA | SB | SI |
| Ed.01 | 0.00 | 3.76 | 1.67 | 4.22 | 0.50 | 0.71 | 2.90 | 2.12 | 28.28 | 28.29 | 29.48 | 28.31 | 28.87 | 28.29 | 28.32 | 22.11 | 22.12 | 21.96 | 21.92 | 22.16 | 30.66 | 30.80 | 28.97 |
| Ed.02 | 3.76 | 0.00 | 3.69 | 5.45 | 3.72 | 3.72 | 4.18 | 4.42 | 28.64 | 28.65 | 29.57 | 28.67 | 29.21 | 28.66 | 28.69 | 22.22 | 22.42 | 22.12 | 22.05 | 22.33 | 30.37 | 30.56 | 28.56 |
| Ed.03 | 1.67 | 3.69 | 0.00 | 4.37 | 1.59 | 1.81 | 2.90 | 2.63 | 28.32 | 28.33 | 29.46 | 28.35 | 28.90 | 28.34 | 28.38 | 22.11 | 22.15 | 21.99 | 21.94 | 22.18 | 30.67 | 30.82 | 29.01 |
| Ed.07 | 4.22 | 5.45 | 4.37 | 0.00 | 4.18 | 4.18 | 5.41 | 4.74 | 28.63 | 28.64 | 29.65 | 28.66 | 29.12 | 28.63 | 28.69 | 22.30 | 22.43 | 22.25 | 22.14 | 22.37 | 30.54 | 30.75 | 28.86 |
| Ed.08 | 0.50 | 3.72 | 1.59 | 4.18 | 0.00 | 0.50 | 2.84 | 2.05 | 28.28 | 28.29 | 29.47 | 28.30 | 28.87 | 28.28 | 28.32 | 22.10 | 22.12 | 21.96 | 21.91 | 22.16 | 30.65 | 30.80 | 28.97 |
| Ed.09 | 0.71 | 3.72 | 1.81 | 4.18 | 0.50 | 0.00 | 2.79 | 1.99 | 28.27 | 28.28 | 29.47 | 28.30 | 28.86 | 28.28 | 28.31 | 22.12 | 22.14 | 21.98 | 21.93 | 22.17 | 30.67 | 30.81 | 28.97 |
| Ed.12 | 2.90 | 4.18 | 2.90 | 5.41 | 2.84 | 2.79 | 0.00 | 3.44 | 28.67 | 28.68 | 29.87 | 28.70 | 28.96 | 28.67 | 28.72 | 22.25 | 22.27 | 22.15 | 22.11 | 22.44 | 30.90 | 31.07 | 29.09 |
| Ed.21 | 2.12 | 4.42 | 2.63 | 4.74 | 2.05 | 1.99 | 3.44 | 0.00 | 28.32 | 28.32 | 29.60 | 28.35 | 28.87 | 28.31 | 28.35 | 22.32 | 22.34 | 22.18 | 22.13 | 22.36 | 30.51 | 30.66 | 28.74 |
| Ed.06 | 28.28 | 28.64 | 28.32 | 28.63 | 28.28 | 28.27 | 28.67 | 28.32 | 0.00 | 1.73 | 8.67 | 1.00 | 4.43 | 1.50 | 1.67 | 28.16 | 28.23 | 27.95 | 27.91 | 28.15 | 32.61 | 32.61 | 31.51 |
| Ed.11a | 28.29 | 28.65 | 28.33 | 28.64 | 28.29 | 28.28 | 28.68 | 28.32 | 1.73 | 0.00 | 8.82 | 0.00 | 4.43 | 0.50 | 1.33 | 28.20 | 28.26 | 27.99 | 27.96 | 28.21 | 32.61 | 32.60 | 31.50 |
| Ed.11b | 29.48 | 29.57 | 29.46 | 29.65 | 29.47 | 29.47 | 29.87 | 29.60 | 8.67 | 8.82 | 0.00 | 8.68 | 9.61 | 8.66 | 9.07 | 29.47 | 29.89 | 29.19 | 29.10 | 29.43 | 32.73 | 32.84 | 31.43 |
| Ed.14 | 28.31 | 28.67 | 28.35 | 28.66 | 28.30 | 28.30 | 28.70 | 28.35 | 1.00 | 0.00 | 8.68 | 0.00 | 4.43 | 1.00 | 1.67 | 28.20 | 28.27 | 27.99 | 27.96 | 28.21 | 32.63 | 32.63 | 31.53 |
| Ed.17 | 28.87 | 29.21 | 28.90 | 29.12 | 28.87 | 28.86 | 28.96 | 28.87 | 4.43 | 4.43 | 9.61 | 4.43 | 0.00 | 4.59 | 4.86 | 28.72 | 28.87 | 28.46 | 28.47 | 28.72 | 32.71 | 32.73 | 31.40 |
| Ed.18 | 28.29 | 28.66 | 28.34 | 28.63 | 28.28 | 28.28 | 28.67 | 28.31 | 1.50 | 0.50 | 8.66 | 1.00 | 4.59 | 0.00 | 1.24 | 28.17 | 28.23 | 27.97 | 27.93 | 28.18 | 32.63 | 32.61 | 31.50 |
| Ed.19 | 28.32 | 28.69 | 28.38 | 28.69 | 28.32 | 28.31 | 28.72 | 28.35 | 1.67 | 1.33 | 9.07 | 1.67 | 4.86 | 1.24 | 0.00 | 28.12 | 28.13 | 27.97 | 27.93 | 28.17 | 32.70 | 32.63 | 31.58 |
| Ed.10 | 22.11 | 22.22 | 22.11 | 22.30 | 22.10 | 22.12 | 22.25 | 22.32 | 28.16 | 28.20 | 29.47 | 28.20 | 28.72 | 28.17 | 28.12 | 0.00 | 4.15 | 3.66 | 3.14 | 4.43 | 30.51 | 30.69 | 26.88 |
| Ed.20 | 22.12 | 22.42 | 22.15 | 22.43 | 22.12 | 22.14 | 22.27 | 22.34 | 28.23 | 28.26 | 29.89 | 28.27 | 28.87 | 28.23 | 28.13 | 4.15 | 0.00 | 4.37 | 4.04 | 5.03 | 30.40 | 30.56 | 26.84 |
| Ed.22 | 21.96 | 22.12 | 21.99 | 22.25 | 21.96 | 21.98 | 22.15 | 22.18 | 27.95 | 27.99 | 29.19 | 27.99 | 28.46 | 27.97 | 27.97 | 3.66 | 4.37 | 0.00 | 2.09 | 3.88 | 30.22 | 30.45 | 26.68 |
| SC | 21.92 | 22.05 | 21.94 | 22.14 | 21.91 | 21.93 | 22.11 | 22.13 | 27.91 | 27.96 | 29.10 | 27.96 | 28.47 | 27.93 | 27.93 | 3.14 | 4.04 | 2.09 | 0.00 | 3.47 | 30.22 | 30.43 | 26.67 |
| SII | 22.16 | 22.33 | 22.18 | 22.37 | 22.16 | 22.17 | 22.44 | 22.36 | 28.15 | 28.21 | 29.43 | 28.21 | 28.72 | 28.18 | 28.17 | 4.43 | 5.03 | 3.88 | 3.47 | 0.00 | 29.69 | 29.88 | 26.27 |
